# Metabolic responses in opposite sun-exposed Antarctic cryptoendolithic communities

**DOI:** 10.1101/725663

**Authors:** Claudia Coleine, Federica Gevi, Giuseppina Fanelli, Silvano Onofri, Anna Maria Timperio, Laura Selbmann

**Affiliations:** Department of Ecological and Biological Sciences, University of Tuscia, Viterbo 01100 Italy; Department of Science and Technology for Agriculture, Forestry, Nature and Energy, University of Tuscia, Viterbo 01100 Italy; Italian National Antarctic Museum (MNA), Mycological Section, Genoa, Italy

**Keywords:** Antarctic cryptoendolithic communities, Sun-exposure, Metabolic activity, Stress-response, Melanin, Allantoin.

## Abstract

Antarctic cryptoendolithic communities are self-supporting borderline ecosystems spreading across the extreme conditions of the Antarctic desert and represent the most predominant life-form in the ice-free desert of McMurdo Dry Valleys, accounted as the closest terrestrial Martian analogue. Components of these communities are very adapted extremophiles and extreme-tolerant microorganisms, among the most resistant known to date. Recent advances started to investigate the biodiversity and community composition in these microbial ecosystems but the metabolic activity of the metacommunity has never been investigated to date. In this study, we explored the stress-response, spreading in two different sites of the same location, subjected to increasing environmental pressure due to opposite sun exposure, to verify the effect of sunlight on settlement and adaptation strategies. Results indicated that the metabolic responses are shaped according to external conditions; in the overall 252 altered metabolites (56 and 196 unique for north and south, respectively), distinguished the two differently exposed communities. We also selected 10 metabolites and performed two-stage Receiver Operating Characteristic (ROC) analysis to test them as potential biomarkers. We focused further on melanin and allantoin as protective substances; their concentration was highly different in the community in the shadow or in the sun.

## INTRODUCTION

The Antarctic cryptoendolithic communities are microbial ecosystems that dominate the biology of the ice-free areas in Continental Antarctica. They were described for the first time in the McMurdo Dry Valleys, or Ross Desert, located in Southern Victoria Land (Friedmann, 1982), the largest among ice-free regions of the continent. The McMurdo Dry Valleys represent a nearly pristine environment largely undisturbed and uncontaminated by humans and show remarkable peculiarities, representing an important analogue for the conditions of ancient Earth and contemporary Mars. They are, in fact, among the most investigated areas as a model environment for astrobiological studies (Horowitz et al., 1972; Wynn-Williams and Edwards, 2000; Onofri et al., 2004). These ice-free areas are dominated mostly by oligotrophic mineral soil and rocky outcrops (Thomas, 1997; Ugolini & Bockheim, 2008) and, for the harsh conditions of low temperatures, always below the freezing point (Doran et al., 2002), strong and rapid thermal fluctuations, extreme dryness, oligotrophy and high UV radiation the biology is dominated by cryptic microbial life-forms dwelling inside rocks (Nienow and Friedmann, 1993). For their unique geological and biological peculiarities, the McMurdo Dry Valleys, as a whole, are now designated as an ASMA (Antarctic Specially Managed Area), to assist planning and coordination of activities, improving cooperation between parties and minimise environmental impacts (SCAR, 2004). The McMurdo Dry Valleys also include five different ASPA (Antarctic Specially Protected Areas) to protect outstanding environments; each ASPA has its own management plan and a specific permit is required for entry. The presence of unusual microhabitats and biological communities, among which cryptoendolithic colonization dominates, are the main reason for requiring the special protection of these areas. There, the prohibitive conditions are incompatible with active life on rock surfaces, and endolithic adaptation confers to microbes the chance to exploit a more protected niche inside rock porosity characterized by a milder and more stable microclimate, where microbes can find refuge in cryptic niches as last chance for survival (Friedmann and Koriem, 1989; Kappen, 1993; Wierzchos and Ascaso, 2001). Antarctic microbial endoliths are among the most stress-resistant organisms known to date, living close to the edge of their physiological adaptability. This life-style must, therefore, be regarded as a border adaptation of life before extinction (Friedmann and Ocampo-Friedmann, 1976; Nienow and Friedmann, 1993).

Due to their high degree of adaptation, these communities are very sensitive to any external perturbation (Selbmann et al., 2017; Coleine et al., 2018a,b); the effect of global warming is dramatically intense at the Poles (Bekryaev et al., 2010) and disturbance due to the effect of Climate Change may have irreversible effects on the weak equilibrium of these communities (Selbmann et al., 2012; 2017). Thus, improving our knowledge on functional capacity and multiple stress-responses of these border ecosystems will give us important clues for monitoring and predicting any future variation (Hogg and Wall, 2011; Selbmann et al., 2017). Recent studies have focused on investigate the biodiversity of the McMurdo Dry Valleys (de la Torre et al., 2003; Cary et al., 2010; Wei et al., 2016; Archer et al., 2017; Lee et al., 2019), while others provided new insights on how the combination of environmental parameters of altitude and sea distance influence the distribution and the settlement of cryptoendolithic colonization (Selbmann et al., 2017; Coleine et al., 2018a), and the effect of the sun exposure parameter in shaping the composition and abundance of functional groups of fungi in the Antarctic endolithic communities (Selbmann et al., 2017; Coleine et al., 2018b; 2019 a).

It was previously hypothesized that the distribution of endoliths reflects the degree of insolation on the rock surfaces; in northern exposed rocks, presenting considerably more favourable conditions than southern exposures face, cryptoendolithic colonization is usually observed (Friedmann, 1977; Friedmann and Weed, 1987). Indeed, the capability to maintain biological activity depends on sufficient insolation on the rock surfaces allowing an efficient photosynthetic process; moreover, temperature became warmer and allows metabolic activity for the increasing of aw as consequence of melting of snow (McKay and Friedmann, 1984). Therefore, insolation is doubtless one of the main factors allowing the growth and survival of any microbe (Deegenaars and Watson, 1998): southern exposed faces may be too shady and cold for biological activity.

A key to understanding the adaptation of these communities is exploring the metabolic activity in the natural environments and production of enzymes that are active at low temperatures, allowing them to survive in such as prohibitive conditions.

Antarctic cryptoendolithic microorganisms have to face several stresses simultaneously and they need to concurrently develop survival strategies to address external conditions. A conspicuous number of studies have been performed on the adaptation strategies on black fungi from these communities, resulted highly resistant in terms of extremes of temperatures, acidity, osmotic stress and salinity, dehydration and irradiation (Sterflinger, 2006; Sterflinger et al., 2012; Gorbushina et al., 2003; Dadachova et al., 2007; Onofri et al., 2007; 2008; Selbmann et al., 2017); besides, the stress-response mechanisms of whole metacommunity has not yet been investigated, either on metagenomic or metabolomic level.

In this investigation, for the first time to our knowledge, metabolomics was successfully applied to Antarctic cryptoendolithic communities to start defining adaptation strategies adopted by these communities to survive in these harsh conditions.

We found, different microbes’ response to sun exposure by modulating specific pathways; in particular, among the candidates to be considered biomarkers, the precursor metabolites of melanin and allantoin pathways were the most affected by sun exposure and we may consider these pathways to be directly related to environmental pressure.

## MATERIALS AND METHODS

### Samples collection

Sandstones were collected at Finger Mt. (1720 m a.s.l., McMurdo Dry Valleys, Southern Victoria Land, Continental Antarctica) both from north (77°45’0.93"S 160°44’45.2"E) and south (77°45’10"S 160°44’44.39.7"E) sun-exposed surfaces by Laura Selbmann during the XXXI Italian Antarctic Expedition (Dec. 2015-Jan. 2016) (Fig.1).

**Figure 1.**
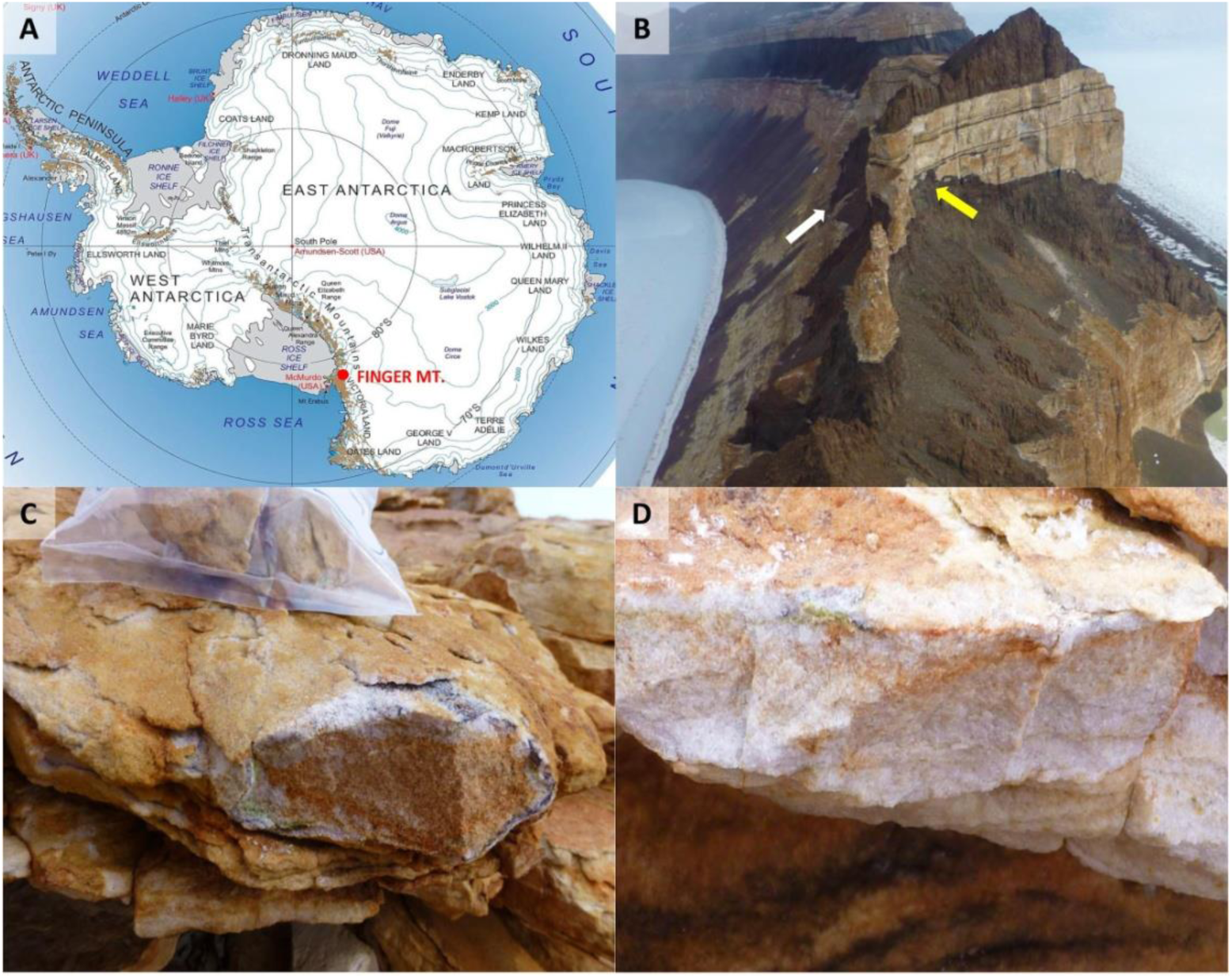
A) Map of Antarctica. Red symbol indicates the study area, Finger Mt. (McMurdo Dry Valleys, Southern Victoria Land, Continental Antarctica). B) Finger Mt. landscape: the yellow arrow indicates the north sun-exposed surface, while the black arrow indicates the sun-exposed surface; north sun-exposed sandstone rock; south sun-exposed sandstone rock.

The presence of endolithic colonization was assessed by direct observation *in situ* using magnification lenses. Rocks were excised using a geological hammer, placed in sterile bags and shipped at −20°C at the at University of Tuscia (Italy) where have been preserved at −20°C in the Mycological Section of the Italian Antarctic National Museum (MNA), until molecular analysis.

### Metabolites extraction

One gram of each rock sample was added to 3000 μl of a chloroform/methanol/water (1:3:1 ratio) solvent mixture stored at −20 °C. Briefly, samples were vortexed for 5 min and left on ice for 2 h for total protein precipitation. Solutions were then centrifuged for 15 min at 15,000×g. and were dried to obtain visible pellets. Finally, the dried samples were re-suspended in 0.1 mL of RNAse/DNAse free water, 5% formic acid and transferred to glass autosampler vials for LC/MS analysis. Extraction was performed in triplicate.

### UHPLC-MS

Twenty-microliter of each extracted supernatant samples were injected into an ultra-high-performance liquid chromatography (UHPLC) system (Ultimate 3000, Thermo) and run on a positive mode: samples were loaded onto a Reprosil C18 column (2.0mm× 150 mm, 2.5 μm-DrMaisch, Germany) for metabolite separation. Chromatographic separations were made a flow rate of 0.2 ml/min. A 0–100% linear gradient of solvent A (ddH2O, 0.1% formic acid) to B (acetonitrile, 0.1%formic acid) was employed over 20 min, returning to 100% A in 2 min and holding solvent A for a 6-min post time hold. The UHPLC system was coupled online with a Q Exactive mass spectrometer (Thermo Fisher, Rockford, IL) scanning in full MS mode (2 μ scans) at resolution of 70,000 in the 67 to 1,000 m/z range, with 3.8 kV spray voltage, 40 sheath gas, and 25 auxiliary gas. The system was operated in positive ion mode. Calibration was performed before each analysis against positive or negative ion mode calibration mixes (Thermo Fisher) to ensure error of the intact mass within the sub ppm range. Metabolite assignments were performed using MAVEN v5.2 (Clasquin et al., 2012) Each replicate was analysed separately and a p-value < 0.01 was tested for all abundance comparisons between sets of triplicates.

### Data elaboration and statistical analysis

Replicate files were processed through MAVEN v5.2, enabling rapid and reliable metabolite quantitation from multiple reaction-monitoring data or high-resolution full-scan mass spectrometry data. Mass Spectrometry chromatograms were created for peak alignment, matching and comparison of parent and fragment ions with tentative metabolite identification within a 2-p.p.m. mass-deviation range between the observed and expected results against an imported KEGG database (Kanehisa et al., 2000).

To visualize the number of significantly changed m/z values between Finger Mt. north and Finger MT. south, a Volcano plots were created using the MetaboAnalyst 3.0 software (http://metpa.metabolomics.ca/). Raw data were normalized by sum and auto-scaling in order to increase the importance of low-abundance ions without significant amplification of noise. This type of plot displays the fold change differences and the statistical significance for each variable. The log of the fold change is plotted on the x-axis so that changes in both directions (up and down) appear equidistant from the centre. The y-axis displays the negative log of the p-value from a two-sample t-test. False discovery rate (FDR) (Benjamini, and Hochberg, 1995) were used for controlling multiple testing.

Metabolites subject to major changes were displayed and Receiver Operator Characteristic (ROC) curves for each of them was calculated using the same software to evaluate potential metabolites to be considered as biomarkers. A ROC diagram plots the true positive rate (sensitivity) of a test on the y-axis against the false positive rate (100-specificity) on the x-axis, yielding the ROC area under the curve (AUC). Within this model, the ROC curves used 5-fold cross validation. AUC can be interpreted as the probability that a test or a classifier will rank a randomly chosen positive instance higher than a randomly chosen negative one. If all positive samples are ranked before negative ones (i.e. a perfect classifier), the AUC is 1.0. An AUC of 0.5 is equivalent to randomly classifying subjects as either positive or negative (i.e. the classifier is of no practical utility).

Metabolic pathways were displayed with Graphpad Prism v5.01 (Graphpad Software Inc) and statistical analyses were performed with the same software (p < 0.05). Data are presented as mean ± SD. Differences were considered statistically significant at *p < 0.05 and further stratified to **p < 0.01, respectively.

## RESULTS

### Metabolites extraction

An untargeted metabolomics analysis was performed on differently sun-exposed rocks and more than 10,000 peaks per sample were referring to the KEGG database; among them, 2807 metabolites were analysed more precisely and identified.

The significant discriminating metabolites were identified using the Volcano plot analysis (Fig. 2). The univariate analysis identified significant accumulation of specific metabolites; most of them were expressed in rocks collected in south-exposure, while few others were overexpressed in north-exposed rocks.

**Figure 2.**
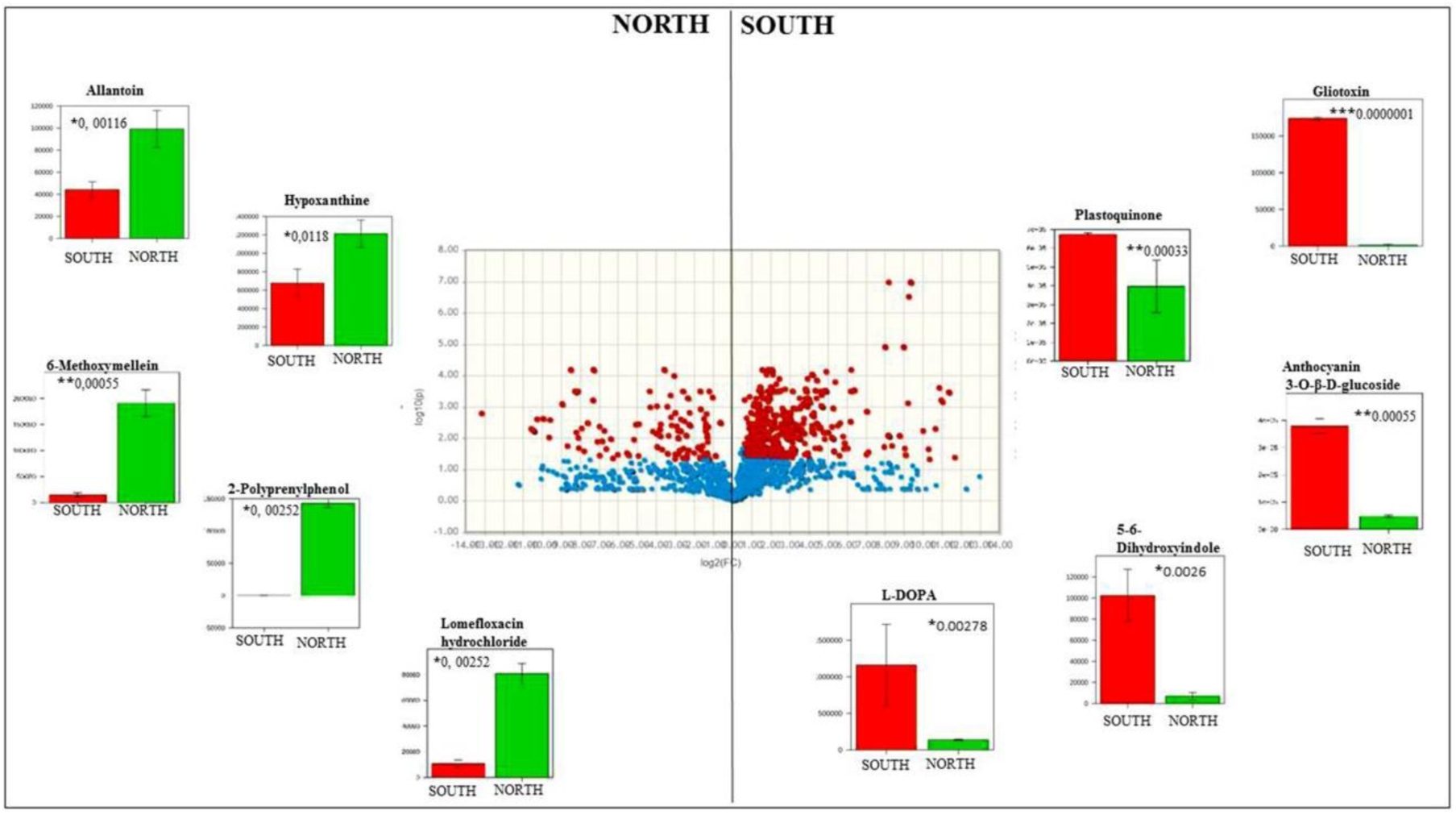
Volcano plot analysis of metabolic changes Finger Mt. north and Finger MT. south. Each point on the volcano plot was based on both p-value and fold-change values, and in this study these two values were set at 0.05 and 2.0, respectively. The points which satisfy the condition p < 0.05 and fold change > 2.0 appear in red and are marker candidates, whereas the others appear in blue that are not significant. The potential biomarkers between experimental groups were annotated with their matched metabolite names. Bar plots show the original values (mean +/-SD). Differences were considered statistically significant at *p < 0.05 further stratified to **p < 0.01 and ***p < 0.001 respectively

Based on the specific criteria, 56 metabolites were significantly upregulated in Antarctic cryptoendolithic communities of rocks exposed to the north whereas 196, were significantly upregulated in the south exposed ones. Supplementary Tables S1 summarized all metabolites identified considering the following parameters: p value, FC, log2(FC), −log10(p).

Based on the available literature, se selected some metabolites to test the performance of the optimal model, using a Receiver Operating Characteristic (ROC) curve analysis and the validation data set. Five metabolites significantly increased in north exposed (Fig. 2) (*Allantoin, Hypoxanthine, 6-Methoxymellein, 2-Polyprenylphenol* and *lomefloxacin hydrochloride*) and five metabolites significantly increased in south exposed (Fig. 3) (Gliotoxin, Plastoquinone, Anthocyanin 3-O-β-D-glucoside, 5-6 Dihydroxyindole and L-DOPA) have been selected to test if they can be biomarkers, associated to a specific environmental condition (i.e. north or south sun-exposure) (see Fig. 2). ROC analysis using this set of 10 metabolites yielded an AUC = 1 as shown in Figs. 3 and 4, which represented a perfect ROC curve according to the accepted classification of biomarkers (Andersen et al., 2014). Among the metabolites considered for ROC analysis, changes in Allantoin (northern rock surface) and L-DOPA (southern rock surface) expression were investigated for their key roles in the Antarctic cryptoendolithic communities.

**Figure 3.**
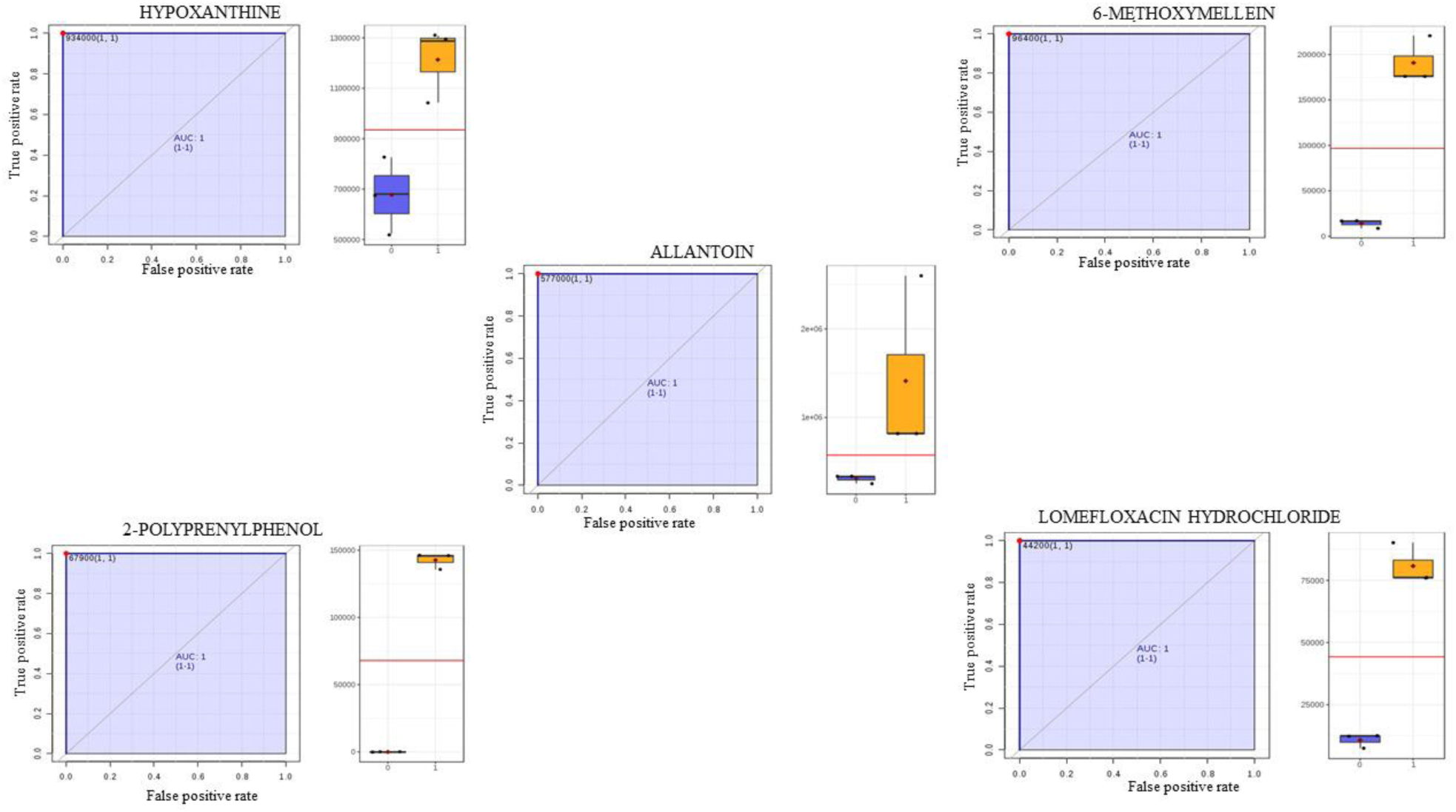
Area under the receiver-operating characteristic curves (AUROC) of the significant metabolite (with AUROC value>0.9) (left panel) with their respective boxplots (right panel) identified 5 metabolites (Allantoin, Hypoxanthine, 6-Methoxymellein, 2-Polyprenylphenol and Lomefloxacin hydrochloride) as significantly elevated in Finger Mt. north samples. They are candidate biomarkers with excellent value (AUC = 1; CI = 1−1).

**Figure 4.**
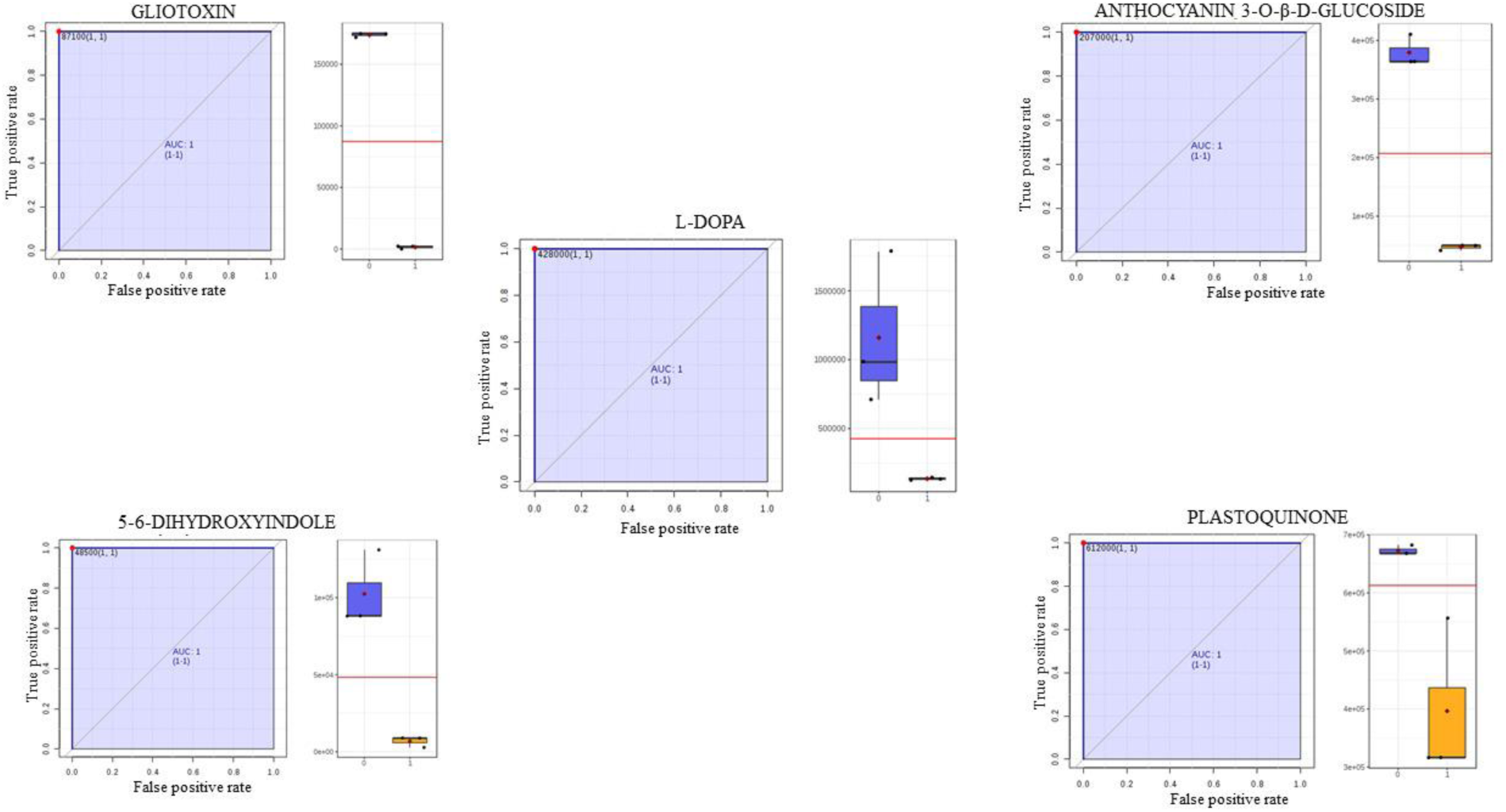
Area under the receiver-operating characteristic curves (AUROC) of the significant metabolite (with AUROC value>0.9) (left panel) with their respective boxplots (right panel) identified 5 metabolites (Gliotoxin, Plastoquinone, Anthocyanin 3-O-β-D-glucoside, 5-6 Dihydroxyindole and L-DOPA) as significantly elevated in Finger Mt. south samples. They are candidate biomarkers with excellent value (AUC = 1; CI = 1−1).

### Allantoin pathway

Figure 5 shows the significant changes observed in the intermediates allantoin biosynthesis. Biosynthesis starts from purine metabolism degradation (Hypoxanthine and Xanthine) and the intermediate Uric Acid. All allantoin pathway intermediates were stable and more abundant in the north exposure; while, an intermediates fluctuation was observed in the south-samples. The amount of uric acid and allantoin decrease significantly in south-exposed samples (p < 0.05).

**Figure 5.**
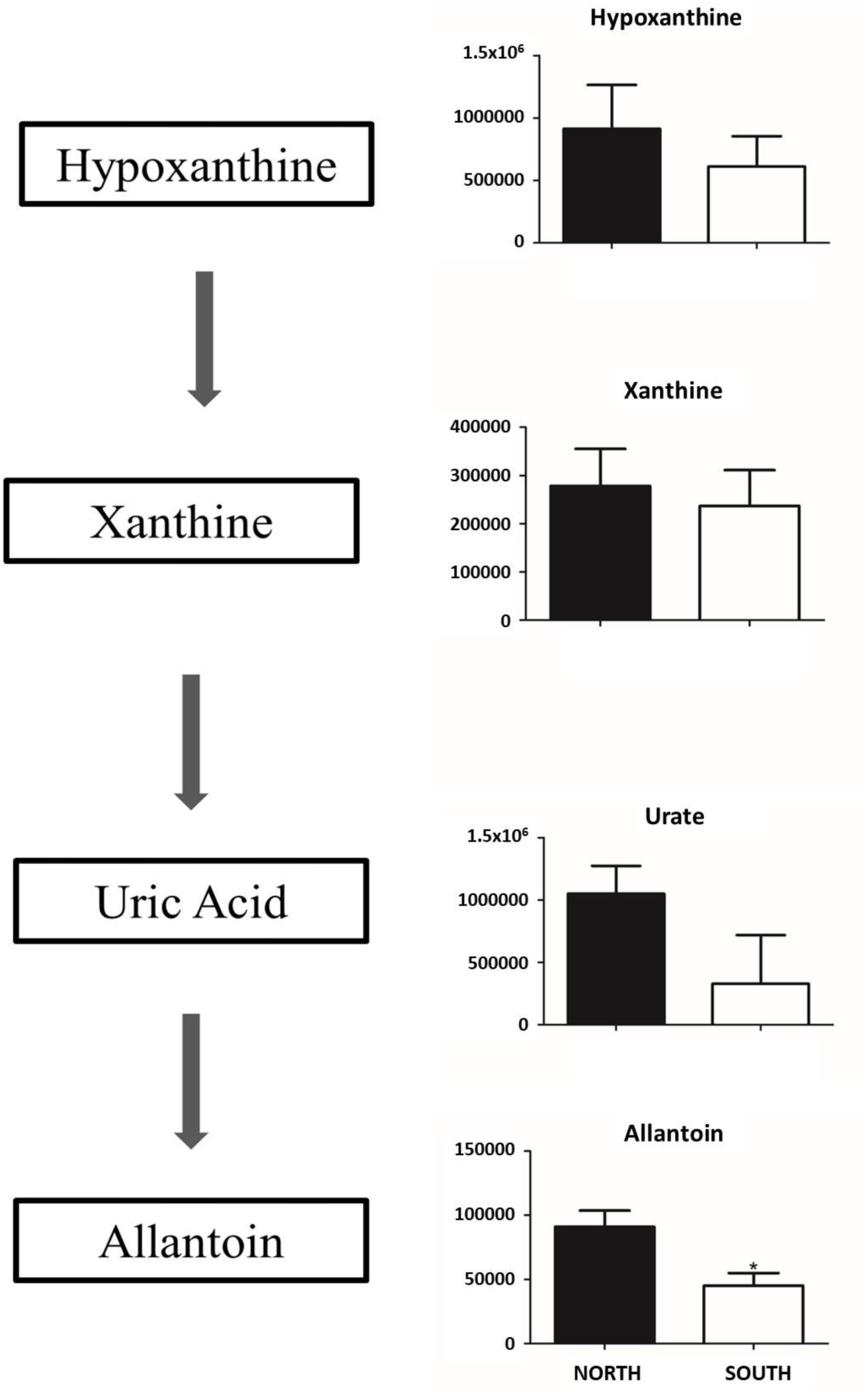
Allantoin pathway. Total amount of allantoin pathway intermediates in the Sandstones appears to be increased in Finger Mt. North with respect to Finger Mt. south. Metabolites are expressed as the mean ± SD concentration over Finger Mt. north *p < 0.05 against Finger Mt. south.

### Melanin pathway

In Fig. 6 the melanin biosynthetic pathway is reported. Melanin is produced starting from oxidation of amino acid L-Tyrosine. With our mass spectrometer techniques, it is possible to identify the tyrosine metabolism intermediates. The rate-limiting initial step in the biosynthesis of melanin is the hydroxylation of tyrosine to L-3,4-dihydroxyphenylalanine (DOPA) and its immediate subsequent oxidation to DOPAquinone (DQ). Dopaquinone went through instantaneous intramolecular nonenzymatic cyclization forming leucochrome, which is rapidly oxidized by dopaquinone to dopachrome at the level of DOPAquinone point of switch. The pathway lead to the synthesis of melanin instead of cysteinyl DOPA and all the intermediates leucoDOPAchrome, DOPAchrome, 5-6 dihydroxyindole, Indole 5,6 quinone reach the maximum level in south sun-exposed samples (p< 0.05).

**Figure 6.**
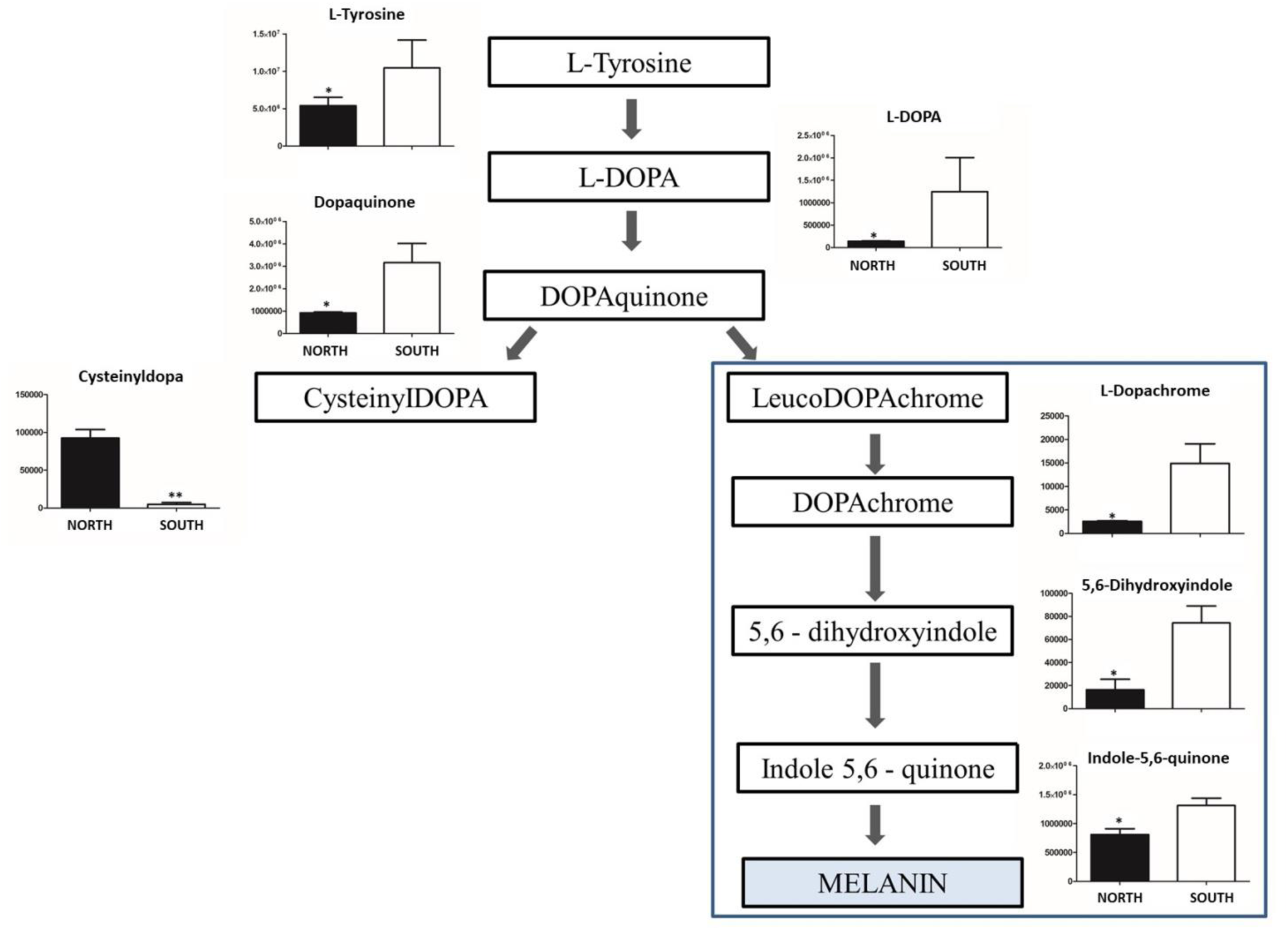
Melanin synthesis pathway. The conversion of tyrosine to melanin involves several steps and we found overall up-regulation of Finger Mt. south intermediates whereas CysteinyIDOPA decrease, confirming that samples from south increase the synthesis of melanin. Metabolites are expressed as the mean ± SD concentration over Finger Mt. north *p < 0.05, **p < 0.01 against Finger Mt. south.

## DISCUSSION

Understanding the mechanisms driving functional and ecological processes in the extremes remains a major challenge (Casanueva et al. 2010; Andrei et al. 2012; Makhalanyane et al. 2015), particularly since environmental stressors are often associated with diminished ecosystem capacity and functionality (Petchey et al. 1999; Schimel et al. 2007; Harrison et al. 2013; Ferrenberg et al. 2015). This is particularly true in hot and cold desert ecosystems; among them the ice-free areas of McMurdo Dry Valleys (Victoria Land, Continental Antarctica) are accounted as the coldest and most hyper-arid desert on Earth and the closest terrestrial Martian analogue (Doran *et al*., 2010; Tamppari *et al*., 2012). The main goal of our study was to understand the responses to different degree of stress of Antarctic cryptoendolithic communities in this area, highlighting the differences in metabolic profiles between communities of north or south exposed rock faces. Metabolites are the result of both biological and environmental factors, and provide great potential to bridge knowledge of genotype and phenotype. The potential of a metabolomics method to detect statistically significant perturbations in the metabolome of an organism is enhanced by excellent analytical precision, unequivocal identification and broad metabolomic coverage. In contrast to targeted metabolomics, non-targeted approaches offer the potential to determine biomarkers. The idea that untargeted mass spectrometry (MS)-based metabolomics analysis will result in a large list of ‘identified’ small molecules that can be mapped to networks and pathways is often assumed (Schrimpe-Rutledge et al., 2016).

In this study, we considered rocks collected at opposite sun-exposure, accounted among the principal driving factor for settlement and shaping stress-response in cryptoendolithic life-style; indeed, recently, sun-exposure was found to be critical in shifting community composition, taxon abundance, and distribution of functional groups of fungi in Antarctic endolithic communities (Coleine *et al*., 2018b; Coleine *et al*., 2019a), affecting these ecosystems more than other abiotic parameters considered (e.g. altitude). Moreover, it was previously hypothesized that the distribution of endoliths reflects the degree of insolation on the rock surfaces; in northern exposed rocks cryptoendolithic colonization is more often observed than southern exposed faces (Friedmann, 1977; Friedmann and Weed, 1987) because warmer temperatures leads to higher liquid water availability to sustain the metabolic activity (McKay and Friedmann, 1985; Deegenaars and Watson, 1998).

Our results clearly showed that the sun exposure strongly influences the metabolic activity in the communities. Indeed, through the Volcano Plots, we highlighted the differential abundance of metabolites influenced by this parameter.

In total, we identified 56 metabolites that were significantly expressed in north sun-exposed rocks and 196 in southern samples as showed in Fig. 2 (p < 0.01, Supplementary Table S1).

The concentration and composition of metabolites presented a clear pattern of correlation across different sun-exposed surfaces of the same mountain, where the changes in the microbial community metabolic profiles following sunlight deprivation were evident. Overall, the remarkable variability observed across sun-exposures indicated that this environmental parameter can be considered as driving factor shaping the stress-response strategies in the Antarctic cryptoendolithic communities. Additionally, the volcano plot analysis, coupled with Receiver Operating Characteristics (ROC) curves analysis, was employed to explore the chemical markers that contributed most to the difference in chemical profiles across sampled area, determining which compounds were of interest. Among the detected metabolites, based on available literature, we used volcano plots to produce a hit of metabolites specimens and identified at least 10 metabolites that were specifically correlated to a specific conditions. Biomarkers are commonly defined as measured characteristics that may be used as indicators of some biological state or condition; an area under the curve (AUC) is a measure of the accuracy of a biomarker test: a value of 1.0 indicates a perfect test (due to absence of overlap of the test data from the two analysed conditions), whereas an AUC value of 0.5 shows the test is no better than random chance. The ROC curves (Figs. 3-4), determined that *allantoin, hypoxanthine, 6-methoxymellein, 2-polyprenylphenol*, and *lomefloxacin hydrochloride* were the most sensitive and specific biomarkers for north sun-exposed rocks (AUC = 1), while *l-dopa, 5-6-dihydroxindole, gliotoxin, plastoquinone*, and *anthocyanin* were unique for south sun-exposed microbial communities (AUC = 1). All these metabolites are intermediates of crucial pathways responsible for stress-adaptation: among the north-related biomarkers, *6^−^methoxymellein* has been reported to be involved in UV-radiation response in plant systems (Mercier et al., 1994); *2-polyprenylphenol* (member of the class of phenols and precursor of ubiquinone) plays an important role in plant resistance and in plant growth and development, through the biosynthesis compounds that act as antioxidants (Liu and Lu, 2016). Their synthesis is generally triggered in response to biotic/abiotic stresses and especially under salt stress conditions; they take part in defence responses during infection, excessive sun exposure, injuries and heavy metal stress (Souza and Devaraj, 2010).

Instead, among biomarkers responsible for stress-response in south-exposed communities, *gliotoxin* is involved in protecting *A. fumigatus* against oxidative stress against ROS (Scharf et al., 2010; Schrettl et al., 2010); indeed, *gliotoxin* is an epipolythiodioxopiperazine (ETP), a toxic secondary metabolites made only by fungi with molecular mass 326 Da and contains a disulphide bridge which can undergo repeating cleavage and reformation, thereby resulting in a potent intracellular redox activity (Choi, et al, 2007). *Plastoquinone* participates in the metabolism of various important chemical compounds, acting as antioxidants, involved in plant response to salinity stress (Takahashi and Murata, 2008, Suzuki et al., 2012). Finally, *anthocyanins* are a well-known water-soluble pigments found in all plant tissues throughout the plant kingdom, involved in UVB radiation, water stress and particularly cold temperatures (Chalker-Scott, 1999), as reported in *Arabidopsis* (Graham, 1998; Leyva et al., 1995). Indeed, McKown et al. (2013) suggested a sort of commonality between anthocyanin biosynthesis and freezing tolerance, as for *Arabidopsis* mutants deficient in freezing tolerance were unable to accumulate anthocyanins perhaps as a result of increased tolerance of cool temperatures.

Among the 10 metabolites analysed as biomarkers, we focalized our attention on *allantoin* and *hypoxanthine* over expressed in north, and *DOPA* and *5-6 hydroyndole* over expressed in south since they can play a role in different stress responses in the endolithic communities of the two-opposite sun-exposed surfaces. The *allantoin* plays an important protective role to excessive sun exposure; this pathway resulted over-expressed in the north exposed communities, where climatic conditions are more buffered and allow a successful settlement of endolithic colonization. Besides, the degree of insulation is much higher than in the south and photosynthetic organisms such as cyanobacteria and algae need to be protected by UV irradiation. *Allantoin* is also produced in plants, providing protective function by blocking UV-B before it reaches cells, the consequences of UV-B exposure such as repairing damaged DNA (Lois, 1994; Smirnoff, 1998). Furthermore, metabolomics analysis revealed allantoin as a major purine metabolite in *Arabidopsis thaliana* and *Oryza sativa* under various stress conditions such as drought (Oliver et al., 2011; Silvente et al., 2011; Yobi et al., 2013), high salinity (Kanani et al., 2010; Wang et al., 2016), low temperature (Kaplan et al., 2004) and nutrient constraint (Nikiforova et al., 2005; Rose et al., 2012; Coneva et al., 2014), environmental characteristics that are exacerbated in Antarctic ice-free areas.

Conversely, the south sun-exposed surface are in more prohibitive climatic conditions and in the shadow over the entire year, with limited possibility for metabolic activity. In this environment, *melanin* pathway resulted more expressed that in the northern surface. *Melanins* are ancient biological pigments found in all kingdoms of life and in particular, in the fungal kingdom, is present across all phyla with remarkable physicochemical properties, allowing them to perform diverse functions in biological systems (Cordero *et al*., 2017 a, b, c; 2018). Some fungal species are constitutively melanized are referred to as black, dematiaceous, micro colonial or meristematic fungi (Sterflinger et al., 2006) with a worldwide distribution, typically colonizing harsh environmental niches not suitable to most life forms such as as saltpans, hydrocarbon-contaminated sites, exposed bare rocks and monuments, icy habitats, deserts, solar panels and building roofs (Adams et al., 2006; Abdel-Hafez et al., 1994; Friedmann, 1982; Gunde-Cimerman et al., 2000, 2003; Sterflinger et al., 2012; Selbmann et al., 2015; Ruibal et al., 2018); these fungi are also invariably present in Antarctic cryptoednolithic communities (Selbmann et al., 2005, 2008, 2015; Egidi et al., 2014; Coleine et al., 2018 a, b). Interestingly, we found that *melanin* are particularly over-expressed in south faced communities where the conditions are particularly stressing and the need of a super-protective shield if of utmost importance for the survival of the community as a whole. These results are confirmed in a recent study where the authors showed that black fungi were invariably predominant in communities sampled in southern exposed rocks (Coleine et al., 2018b). Black melanized fungi in these communities form a black barrier just above the photobiont’s stratification, supplying protection to algae by dissipating excessive sunlight (Selbmann et al., 2013). The presence of black fungi play, therefore, a crucial role in the survival of the whole community under in these highly stressful conditions characterized by drastic temperature fluctuations, elevated radiation exposure, high osmotic pressure, oxidative stresses, low water activity and nutrient availability and intense sunlight and UV radiation (Dadachova and Casadevall, 2008; Onofri et al., 2008, 2012, 2015, 2018, 2019; Selbmann et al., 2011, 2018).

This first contribution allowed us to investigate the functionality of Antarctic endoliths demonstrating how the metabolic response shifts across variation due to sun exposure.

We validated application of a untargeted metabolomics method that provides optimal identification of a wide range of metabolites to detect metabolic changes in the main pathways, determining which products are being released into the environment.

We conclude that further investigations on changes to the metabolic profiles may have potential for use as early indicators, to forecast the impact of Global Change that is challenging for the structure and functioning of cryptoendolithic ecosystems in Antarctic deserts.

## Acknowledgment

L.S. and C.C. wish to thank the Italian National Program for Antarctic Researches (PNRA) for funding sampling campaigns and researches in Italy in the frame of the PNRA projects. The Italian Antarctic National Museum (MNA) is kindly acknowledged for financial support to the Mycological Section on the MNA for preserving Antarctic rock samples, herein analysed, stored in the Culture Collection of Fungi from Extreme Environments (CCFEE), University of Tuscia, Italy.

## Author contributions

LS, AMT and CC designed research; AMT designed the experiments of mass spectrometry; GF and FG carried out most of the experiments and assisted with the statistical analysis; LS, AMT and CC wrote the manuscript with the inputs of SO, GF and FG.

