## Supplementary Table for "Metabolic responses in opposite sun-exposed Antarctic cryptoendolithic communities"

| **NORTH SUN EXPOSED ROCK SAMPLE** |  |  |  |  |
| --- | --- | --- | --- | --- |
| **Metabolite** | **FC** | **log2(FC)** | **p value** | **-log10(p)** |
| Allantoin | 5.9232 | -2.56 | 0.001133 | 2.95 |
| Hypoxanthine | 0.0657 | -3.92 | 0.011882 | 1.24 |
| Falcarinol | 0.0028 | -8.48 | 6.92E-05 | 4.16 |
| Aricine | 0.0064 | -7.29 | 6.92E-05 | 4.16 |
| 5-Hydroxy-L-tryptophan | 0.0841 | -3.57 | 6.92E-05 | 4.16 |
| Nicotyrine | 0.1204 | -3.05 | 0.000108 | 3.97 |
| 1-Hydroxy-2-aminoethylphosphonate | 0.3102 | -1.69 | 0.000333 | 3.48 |
| Scutellarein | 0.0033 | -8.24 | 0.000334 | 3.48 |
| Thiethylperazine | 0.0037 | -8.10 | 0.000334 | 3.48 |
| Miraxanthin-III | 0.0493 | -4.34 | 0.000334 | 3.48 |
| 2-Oxo-7-methylthioheptanoic acid | 0.4287 | -1.22 | 0.000334 | 3.48 |
| 4-4-Difluoro-17beta-hydroxyandrost-5-en-3-one propionate | 0.1054 | -3.25 | 0.000449 | 3.35 |
| Tauropine | 0.4865 | -1.04 | 0.000523 | 3.28 |
| 6-Methoxymellein | 0.198 | -2.34 | 0.000556 | 3.26 |
| Istamycin B1 | 0.0063 | -7.31 | 0.000647 | 3.19 |
| Broussonin C | 0.0021 | -8.92 | 0.000854 | 3.07 |
| Methyl | 0.0641 | -3.96 | 0.000993 | 3.00 |
| Cyclohexylsulfamate | 0.0936 | -3.42 | 0.001033 | 2.99 |
| Simazine | 0.0503 | -4.31 | 0.001074 | 2.97 |
| alpha-Zearalanol | 0.315 | -1.67 | 0.001246 | 2.90 |
| Sarmentosin | 0.1249 | -3.00 | 0.001626 | 2.79 |
| 20-21-21-Trifluoro-3-methoxy-19-nor-17alpha-pregna | 0.2104 | -2.25 | 0.001626 | 2.79 |
| Tween 80 | 0.0001 | -13.15 | 0.001691 | 2.77 |
| Rutecarpine | 0.3899 | -1.36 | 0.001757 | 2.76 |
| Butoctamide hydrogen succinate | 0.1069 | -3.23 | 0.002254 | 2.65 |
| Lomefloxacin hydrochloride | 0.3548 | -1.50 | 0.002516 | 2.60 |
| 2-Polyprenylphenol | 0.001 | -9.92 | 0.00252 | 2.60 |
| RU 5135 | 0.0008 | -10.25 | 0.002613 | 2.58 |
| Curacin A | 0.1531 | -2.71 | 0.002625 | 2.58 |
| Penicillin K | 0.0013 | -9.56 | 0.002754 | 2.56 |
| Tolbutamide | 0.1754 | -2.51 | 0.00302 | 2.52 |
| Bethanechol | 0.0052 | -7.59 | 0.003458 | 2.46 |
| N-Acetyl-ala-ala-ala-methylester | 0.0333 | -4.91 | 0.00371 | 2.43 |
| 4-4-Disubstituted cyclohexenone | 0.3806 | -1.39 | 0.003716 | 2.43 |
| 2-Acetamidofluorene | 0.0737 | -3.76 | 0.003788 | 2.42 |
| Sertraline | 0.0299 | -5.06 | 0.00386 | 2.41 |
| DL-Pipecolic acid | 0.2099 | -2.25 | 0.003993 | 2.40 |
| Candimine | 0.1098 | -3.19 | 0.004085 | 2.39 |
| Sterol 3-beta-D-glucoside | 0.1324 | -2.92 | 0.004374 | 2.36 |
| 2-Demethylmenaquinone | 0.0080 | -6.97 | 0.004435 | 2.35 |
| thymine | 0.3775 | -1.41 | 0.004541 | 2.34 |
| Carisoprodol | 0.4169 | -1.26 | 0.004541 | 2.34 |
| Sarothralin | 0.0042 | -7.91 | 0.005135 | 2.29 |
| Harzianopyridone | 0.0007 | -10.56 | 0.005297 | 2.28 |
| Nalpha-Benzyloxycarbonyl-L-leucine | 0.0716 | -3.80 | 0.005493 | 2.26 |
| Questinol | 0.0007 | -10.45 | 0.005974 | 2.22 |
| 4-4--DiphenylethenylidenebisN-N-dimethylbenzenamine | 0.0095 | -6.72 | 0.006104 | 2.21 |
| Dibenza-hanthracene | 0.1382 | -2.86 | 0.006125 | 2.21 |
| D-Galactosyl-1-3-beta-D-galactosyl-1-4-beta-D-glucosyl-R | 0.3787 | -1.40 | 0.006218 | 2.21 |
| Tolylacetonitrile | 0.1533 | -2.71 | 0.006253 | 2.20 |
| Atrazine | 0.0555 | -4.17 | 0.006405 | 2.19 |
| Thiamine aldehyde | 0.0034 | -8.19 | 0.006838 | 2.17 |
| 2-Hydroxyiminostilbene | 0.0625 | -4.00 | 0.007567 | 2.12 |
| UH-301 | 0.4011 | -1.32 | 0.008269 | 2.08 |
| Indinavir sulfate | 0.0980 | -3.35 | 0.008747 | 2.06 |
| 4-2-Pyrazinylethenylphenol | 0.0721 | -3.79 | 0.00881 | 2.06 |
| 3-6-8-Trimethylallantoin | 0.3982 | -1.33 | 0.009825 | 2.01 |
| UMP | 0.0013 | -9.62 | 0.009938 | 2.00 |
| **SOUTH SUN EXPOSED ROCK SAMPLE** |  |  |  |  |
| L-DOPA | 22.704 | 4.50 | 0.0278 | 1.56 |
| Dihydroxyindole | 38.948 | 5.29 | 0.00269 | 2.58 |
| Glitoxin | 293.87 | 8.20 | 0.0000 | 6.96 |
| p-Glucosyloxymandelonitrile | 651.36 | 9.35 | 0.0000 | 6.96 |
| 1-6-Dinitropyrene | 257.84 | 8.01 | 0.0000 | 4.89 |
| Scandoside methyl ester | 506.65 | 8.98 | 0.0000 | 4.89 |
| citrulline | 2.9662 | 1.57 | 0.0001 | 4.16 |
| Dihydromethysticin | 3.6798 | 1.88 | 0.0001 | 4.16 |
| 3-Phenylpropyl acetate | 4.1601 | 2.06 | 0.0001 | 4.16 |
| 2-2-Dimethyl-3-4-bis4-methoxyphenyl-2H-1-benzopyran-7-ol acetate | 30.301 | 4.92 | 0.0001 | 4.10 |
| N4-Phosphoagmatine | 3.4328 | 1.78 | 0.0001 | 4.01 |
| Phytosphingosine | 4.3557 | 2.12 | 0.0001 | 4.01 |
| hydroxyphenylpyruvate | 15.333 | 3.94 | 0.0001 | 3.88 |
| 1-Hydroxyalkyl-sn-glycerol | 5.4921 | 2.46 | 0.0001 | 3.87 |
| Pipobroman | 21.518 | 4.43 | 0.0001 | 3.83 |
| 11-Deoxocucurbitacin I | 41.935 | 5.39 | 0.0002 | 3.77 |
| Sulfadoxine | 4.9139 | 2.30 | 0.0002 | 3.71 |
| Chrysanthetriol | 2.7453 | 1.46 | 0.0002 | 3.61 |
| Warburganal | 4.052 | 2.02 | 0.0002 | 3.61 |
| 6-Imino-5-oxocyclohexa-1-3-dienecarboxylate | 1841.3 | 10.85 | 0.0003 | 3.59 |
| Streptobiosamine | 6.8079 | 2.77 | 0.0003 | 3.55 |
| 2-Aminomuconate semialdehyde | 14.296 | 3.84 | 0.0003 | 3.55 |
| Homostachydrine | 84.149 | 6.39 | 0.0003 | 3.51 |
| Huperzine B | 2.6191 | 1.39 | 0.0003 | 3.50 |
| Loperamide | 5.7924 | 2.53 | 0.0003 | 3.50 |
| Piperidine | 14.745 | 3.88 | 0.0003 | 3.50 |
| N-Acetylneuraminate | 20.403 | 4.35 | 0.0003 | 3.50 |
| Stachyose | 29.427 | 4.88 | 0.0003 | 3.50 |
| Gla protein precursor | 2.7942 | 1.48 | 0.0003 | 3.48 |
| N-Cyclopropylammelide | 3.5946 | 1.85 | 0.0003 | 3.48 |
| Sarcostin | 3.6927 | 1.88 | 0.0003 | 3.48 |
| Plastoquinone | 4.6089 | 2.20 | 0.0003 | 3.48 |
| Coleonol | 5.2495 | 2.39 | 0.0003 | 3.48 |
| Phosphinothricin | 8.9336 | 3.16 | 0.0003 | 3.48 |
| Vicine | 14.51 | 3.86 | 0.0003 | 3.48 |
| Arbutin | 14.519 | 3.86 | 0.0003 | 3.48 |
| L-Serine-phosphoethanolamine | 24.923 | 4.64 | 0.0003 | 3.48 |
| Oxidized Renilla luciferin | 74.086 | 6.21 | 0.0003 | 3.48 |
| Bowdichione | 2618.1 | 11.35 | 0.0004 | 3.44 |
| Dehydrofalcarinol | 4.0094 | 2.00 | 0.0004 | 3.43 |
| Cordycepin | 25.462 | 4.67 | 0.0004 | 3.37 |
| PD 123319 | 2.8655 | 1.52 | 0.0005 | 3.33 |
| Lividomycin B | 11.457 | 3.52 | 0.0006 | 3.26 |
| 6beta-17beta-Dihydroxyandrost-4-en-3-one diacetate | 3.0057 | 1.59 | 0.0006 | 3.26 |
| Buclizine | 3.3833 | 1.76 | 0.0006 | 3.26 |
| 9-Fluoro-11beta-hydroxy-16beta-methylandrosta-1-4-diene-3-17-dione | 3.8093 | 1.93 | 0.0006 | 3.26 |
| Benzenamine sulfate | 3.917 | 1.97 | 0.0006 | 3.26 |
| D-Gal alpha 1-6D-Gal alpha 1-6D-Glucose | 15.453 | 3.95 | 0.0006 | 3.26 |
| Anthocyanin 3--O-beta-D-glucoside | 21.16 | 4.40 | 0.0006 | 3.26 |
| Cycloheximide | 2.3778 | 1.25 | 0.0006 | 3.25 |
| N3--Acetyl-2-deoxystreptamine antibiotic | 14.392 | 3.85 | 0.0006 | 3.25 |
| CDP-choline | 1975.2 | 10.95 | 0.0006 | 3.19 |
| Adifoline | 3.6603 | 1.87 | 0.0007 | 3.14 |
| Neurosporaxanthin | 2084.1 | 11.03 | 0.0007 | 3.13 |
| 7-Methyladenine | 16.63 | 4.06 | 0.0007 | 3.13 |
| 4-Amino-2-hydroxylamino-6-nitrotoluene | 591.8 | 9.21 | 0.0008 | 3.10 |
| Melibiitol | 2.197 | 1.14 | 0.0008 | 3.08 |
| Glycosyl-4-4--diaponeurosporenoate | 2.6635 | 1.41 | 0.0008 | 3.07 |
| Formylanthranilate | 13.191 | 3.72 | 0.0008 | 3.07 |
| Guanidoacetic acid | 2.7054 | 1.44 | 0.0010 | 3.01 |
| Cytochrome c S-methylmethionine | 2.1493 | 1.10 | 0.0010 | 3.00 |
| Nocardicin G | 12.139 | 3.60 | 0.0010 | 3.00 |
| Benzoapyrene-7-8-oxide | 25.455 | 4.67 | 0.0010 | 3.00 |
| Diazinon | 15.334 | 3.94 | 0.0010 | 2.98 |
| 1-Alkyl-2-acylglycerophosphoethanolamine | 9.1331 | 3.19 | 0.0011 | 2.94 |
| Magnocurarine | 38.864 | 5.28 | 0.0012 | 2.94 |
| Evadol hydrochloride | 6.1497 | 2.62 | 0.0012 | 2.93 |
| Chloroprocaine | 2.5138 | 1.33 | 0.0012 | 2.93 |
| Primin | 3.5421 | 1.82 | 0.0012 | 2.93 |
| 7-Acetyloxy-3-3-pyridinyl-2H-1-benzopyran-2-one | 11.732 | 3.55 | 0.0012 | 2.93 |
| N-Isopropylammelide | 10.028 | 3.33 | 0.0012 | 2.92 |
| 5-Ureido-4-imidazole carboxylate | 3.9508 | 1.98 | 0.0012 | 2.91 |
| Flunisolide | 2.6014 | 1.38 | 0.0012 | 2.90 |
| Compound VS | 3.5434 | 1.83 | 0.0013 | 2.90 |
| Amodiaquine | 3.9762 | 1.99 | 0.0013 | 2.87 |
| Indolebutyric acid | 6.3474 | 2.67 | 0.0014 | 2.87 |
| Cinnamaldehyde | 2.6398 | 1.40 | 0.0014 | 2.86 |
| guanine | 6.4786 | 2.70 | 0.0014 | 2.85 |
| Estradiol 17beta-cyclopentylpropionate | 3.2215 | 1.69 | 0.0014 | 2.85 |
| Lasiocarpine | 3.4909 | 1.80 | 0.0014 | 2.84 |
| Tetrahydrozoline | 132.78 | 7.05 | 0.0015 | 2.83 |
| 1-3-4-Dihydroxyphenyl-5-hydroxy-3-decanone | 3.1305 | 1.65 | 0.0016 | 2.81 |
| xanthine | 2.5364 | 1.34 | 0.0016 | 2.79 |
| Phyllanthin | 2.5939 | 1.38 | 0.0016 | 2.79 |
| Guaiazulene | 2.8954 | 1.53 | 0.0016 | 2.79 |
| Istamycin AO | 8.3592 | 3.06 | 0.0016 | 2.79 |
| 7-Hydroxy-6-methyl-8-ribityl lumazine | 19.334 | 4.27 | 0.0016 | 2.79 |
| N6-delta2-Isopentenyl-adenine | 7.7177 | 2.95 | 0.0017 | 2.78 |
| 6-Acetylpicropolin | 2.6915 | 1.43 | 0.0017 | 2.77 |
| N-Methylanthranilamide | 2.3239 | 1.22 | 0.0019 | 2.73 |
| p-hydroxybenzoate | 3.0542 | 1.61 | 0.0019 | 2.71 |
| S-Methyl-3-phospho-1-thio-D-glycerate | 5.0161 | 2.33 | 0.0019 | 2.71 |
| Glycoperine | 6.1411 | 2.62 | 0.0019 | 2.71 |
| Glycerol 1-phosphate | 7.8375 | 2.97 | 0.0019 | 2.71 |
| Pyruvate kinase phosphate | 18.306 | 4.19 | 0.0019 | 2.71 |
| Phenylsulfate | 4.5312 | 2.18 | 0.002 | 2.71 |
| 2S-Amino-tridecanoic acid | 3.3953 | 1.76 | 0.0021 | 2.69 |
| 2-4-Chlorophenyl-3-phenyl-3-2-pyridinylacrylonitrile | 19.24 | 4.27 | 0.0021 | 2.67 |
| S-Substituted N-acetyl-L-cysteine | 2.989 | 1.58 | 0.0022 | 2.65 |
| Methazolamide | 8.0656 | 3.01 | 0.0024 | 2.62 |
| Estrone glucuronide | 3.1205 | 1.64 | 0.0025 | 2.61 |
| BMS-268770 | 13.48 | 3.75 | 0.0025 | 2.61 |
| 1-4-alpha-D-Glucooligosaccharide | 6.6574 | 2.74 | 0.0026 | 2.59 |
| 6--Dehydro-6--oxoparomamine | 8.3311 | 3.06 | 0.0026 | 2.58 |
| 3-Oxo-3-ureidopropanoate | 8.2451 | 3.04 | 0.0027 | 2.57 |
| Gentisyl alcohol | 2.5422 | 1.35 | 0.0027 | 2.56 |
| 2-4-Dihydroxypteridine | 3.3871 | 1.76 | 0.0028 | 2.56 |
| 2-5-Dichloro-4-oxohex-2-enedioate | 4.6216 | 2.21 | 0.003 | 2.52 |
| Dinitrogen reductase | 3.7192 | 1.90 | 0.003 | 2.52 |
| Perazine | 7.6123 | 2.93 | 0.003 | 2.52 |
| S-4-Methylthiobutylthiohydroximoyl-L-cysteine | 2.9828 | 1.58 | 0.0031 | 2.51 |
| Uric acid | 3.3443 | 1.74 | 0.0031 | 2.51 |
| 2-Hydroxy-5-methyl-cis-cis-muconic semialdehyde | 7.5825 | 2.92 | 0.0031 | 2.51 |
| N1-Acetylspermidine | 73.104 | 6.19 | 0.0031 | 2.51 |
| Apoatropine | 2.7146 | 1.44 | 0.0032 | 2.50 |
| Anacyclin | 3.5391 | 1.82 | 0.0033 | 2.48 |
| Thiobinupharidine | 3.253 | 1.70 | 0.0034 | 2.47 |
| Deisopropylhydroxyatrazine | 3.7521 | 1.91 | 0.0034 | 2.47 |
| Metipranolol hydrochloride | 2.6621 | 1.41 | 0.0034 | 2.46 |
| Strigolactone ABC-rings | 2.9212 | 1.55 | 0.0035 | 2.46 |
| 1-1-2-Triphenylpropane | 50.089 | 5.65 | 0.0035 | 2.46 |
| Purine | 3.3104 | 1.73 | 0.0036 | 2.45 |
| N-Acetylphenylethylamine | 3.4987 | 1.81 | 0.0036 | 2.45 |
| WIN IS | 13.135 | 3.72 | 0.0036 | 2.45 |
| Bonafousine | 2.1797 | 1.12 | 0.0036 | 2.45 |
| Columbin | 21.57 | 4.43 | 0.0036 | 2.45 |
| L-2-3-Diaminopropanoate | 4.5345 | 2.18 | 0.0036 | 2.44 |
| Crinamidine | 5.9837 | 2.58 | 0.0036 | 2.44 |
| Benzimidazole | 4.1744 | 2.06 | 0.0037 | 2.43 |
| Kadsurin A | 9.5522 | 3.26 | 0.0037 | 2.43 |
| Pimelea factor P2 | 2.3144 | 1.21 | 0.0037 | 2.43 |
| Bruceine D | 2.8774 | 1.52 | 0.0040 | 2.40 |
| Acanthicifoline | 10.802 | 3.43 | 0.0041 | 2.39 |
| N-Cyclopropylammeline | 18.067 | 4.18 | 0.0041 | 2.38 |
| Eudistomin C | 45.913 | 5.52 | 0.0041 | 2.38 |
| Estradiol-17alpha | 10.121 | 3.34 | 0.0043 | 2.37 |
| Flutamide | 5.8007 | 2.54 | 0.0044 | 2.35 |
| 3-Acetyloxy-9-mercaptoandrosta-3-5-diene-11-17-dione | 6.8368 | 2.77 | 0.0045 | 2.34 |
| Avizafone | 37.212 | 5.22 | 0.0045 | 2.34 |
| Zanamivir | 2.8926 | 1.53 | 0.0048 | 2.32 |
| Clozapine | 4.2078 | 2.07 | 0.0048 | 2.32 |
| Trimethobenzamide | 24.798 | 4.63 | 0.0048 | 2.32 |
| Angustifoline | 2.9005 | 1.54 | 0.0049 | 2.31 |
| Metsulfuron methyl | 5.8017 | 2.54 | 0.0050 | 2.30 |
| D-glucarate | 4.8758 | 2.29 | 0.0050 | 2.30 |
| D-Xylosylprotein | 2.1706 | 1.12 | 0.0051 | 2.29 |
| N-Acetylmuramate | 2.4655 | 1.30 | 0.0051 | 2.29 |
| 1-Ethyl-2-benzimidazolinone | 7.9318 | 2.99 | 0.0051 | 2.29 |
| Furofoline I | 18.228 | 4.19 | 0.0051 | 2.29 |
| trans_trans-farnesyl diphosphate | 6.217 | 2.64 | 0.0051 | 2.29 |
| Pentahomomethionine | 23.302 | 4.54 | 0.0052 | 2.29 |
| LL-2-6-Diaminoheptanedioate | 18.046 | 4.17 | 0.0053 | 2.28 |
| Nepetalactone trans-cis-form | 3.0807 | 1.62 | 0.0053 | 2.28 |
| Acronycidine | 1603 | 10.65 | 0.0053 | 2.28 |
| Abyssinone I | 6.5505 | 2.71 | 0.0053 | 2.28 |
| Butanoylphosphate | 9.5172 | 3.25 | 0.0055 | 2.26 |
| Valproic acid | 7.371 | 2.88 | 0.0055 | 2.26 |
| N3--Acetylkanamycin | 12.411 | 3.63 | 0.0055 | 2.26 |
| Vulgaxanthin-II | 8.3083 | 3.05 | 0.0055 | 2.26 |
| alpha-Amylcinnamaldehyde | 4.0836 | 2.03 | 0.0058 | 2.23 |
| Proacacipetalin | 9.7704 | 3.29 | 0.0061 | 2.21 |
| 2-Hydroxy-6-oxo-6-2-hydroxyphenyl-hexa-2-4-dienoate | 1025 | 10 | 0.0061 | 2.21 |
| Amiloride | 2.2527 | 1.17 | 0.0067 | 2.17 |
| Procollagen trans-4-hydroxy-L-proline | 3.4682 | 1.79 | 0.0067 | 2.17 |
| 2-Butylbenzofuran-3-yl4-hydroxyphenylketone | 18.075 | 4.18 | 0.0067 | 2.17 |
| Deoxyanisatin | 2.7669 | 1.47 | 0.0067 | 2.17 |
| 1-2-Anthracenediol | 11.922 | 3.58 | 0.0068 | 2.17 |
| Valacyclovir | 27.45 | 4.78 | 0.0068 | 2.17 |
| cis-1-2-Dihydro-3-ethylcatechol | 32.201 | 5.01 | 0.0068 | 2.17 |
| gamma-Fagarine | 28.431 | 4.83 | 0.0069 | 2.16 |
| 4-Hydroxymethylphenylhydrazine | 13.63 | 3.77 | 0.0074 | 2.13 |
| Quinidine | 2.6223 | 1.39 | 0.0074 | 2.13 |
| 3-Chloro-L-alanine | 3.6925 | 1.88 | 0.0074 | 2.13 |
| Mechlorethamine | 5.9008 | 2.56 | 0.0074 | 2.13 |
| 2-Polyprenyl-6-hydroxyphenol | 19.257 | 4.27 | 0.0074 | 2.13 |
| Amylopectin | 8.14 | 3.03 | 0.0077 | 2.11 |
| N4-Acetyl-beta-D-glucosaminylasparagine | 8.9674 | 3.16 | 0.0077 | 2.11 |
| 2-Amino-4-oxo-6-1--2--3--trihydroxypropyl-diquinoid-7-8- | 2.7616 | 1.47 | 0.0079 | 2.10 |
| Norcapillene | 3.0381 | 1.60 | 0.0079 | 2.10 |
| 4-Oxocyclohexanecarboxylate | 6.7593 | 2.76 | 0.0079 | 2.10 |
| L-Octanoylcarnitine | 5.6677 | 2.50 | 0.0080 | 2.10 |
| Pseudotropine | 8.3221 | 3.06 | 0.0082 | 2.09 |
| Capsaicin | 7.0086 | 2.81 | 0.0083 | 2.08 |
| Methoxybrassinin | 9.3719 | 3.23 | 0.0087 | 2.06 |
| N-Caffeoylputrescine | 0.187 | -2.42 | 0.0087 | 2.06 |
| Kainic acid | 2.0132 | 1.01 | 0.0087 | 2.06 |
| Diethyl phenyl phosphate | 2.7216 | 1.44 | 0.0087 | 2.06 |
| Toxoflavine | 3.5184 | 1.81 | 0.0087 | 2.06 |
| beta-L-Arabinose 1-phosphate | 289.97 | 8.18 | 0.0087 | 2.06 |
| Alcophosphamide | 435.98 | 8.77 | 0.0088 | 2.06 |
